# Ethmoid bone marrow region modulates lymphoid activation and differentiation with distinct myeloid profile in neuroinflammation

**DOI:** 10.64898/2026.10.03.755016

**Authors:** Collin J Laaker, Kristóf G Kovács, Andy Madrid, Melinda Herbath, Kylee Vi, Jenna Port, Sophia M Vrba, Ajinkya R Limkar, Simon Federico Woen Ordóñez, Skyla Coogan, Thanthrige Thiunuwan Priyathilaka, Martin Hsu, Matyas Sandor, Zsuzsanna Fabry

**Affiliations:** Neuroscience Training Program, University of Wisconsin Madison, Madison, WI, United States; Department of Pathology and Laboratory Medicine, School of Medicine and Public Health, University of Wisconsin Madison, Madison, WI; Cellular and Molecular Pathology Training Program, University of Wisconsin-Madison; Medical Scientist Training Program, University of Wisconsin-Madison, Madison, WI; Cellular and Molecular Pharmacology Training Program, University of Wisconsin-Madison, Madison, WI; Department of Urology, University of Wisconsin-Madison, Madison, WI

## Abstract

The skull bone marrow axis has recently emerged as an important source of immune cells during neuroinflammation, including in the experimental autoimmune encephalomyelitis (EAE), the mouse model of multiple sclerosis. The cribriform plate bone marrow (cpBM), is a portion of the ethmoid bone of the skull of mice, and is positioned next to nasal regions. At the cribriform plate olfactory nerves pass through the skull and recent evidence supports that it represents an important immune niche with links to CSF outflow and connections to cpBM. Previous data supports cellular composition of the ethmoid bone marrow in mice is altered in neuroinflammation suggesting it senses inflammatory cues, but the exact nature of these changes is still unclear. Here using scRNAseq and flow cytometry of the ethmoid bone marrow region we test how the region responds to EAE. In addition, we also compare the cpBM region with systemic bone marrow in the femur to understand region specific differences. We report that neuroinflammation creates a regulatory response at these sites and implicates the cpBM in controlling these effects. A better understanding of the cribriform niche may offer new treatments for neuroinflammatory disease

## Introduction

Multiple sclerosis (MS) and its animal model experimental autoimmune encephalomyelitis (EAE), are characterized by infiltration of peripheral immune cells into the CNS, resulting in demyelination (Rangachari and Kuchroo 2013). While CD4+ T cells are the key pathogenic cell type involved in EAE, studies have also implicated the bone marrow sites as sites of pathogenic pro-inflammatory myeloid cell creation or myelopoiesis (Shi et al. 2022), immune cell priming (Dobersalske et al. 2024) and cell-trafficking into the CNS (Whiteley et al. 2024; Cugurra et al. 2021). Recent advances in skull bone marrow anatomy have uncovered osseous channels between the dura and skull bone marrow that allow immune cell trafficking between regions (Mazzitelli et al. 2023; Cugurra et al. 2021). While systemic bone marrow such as the femur have been largely studied in neuroinflammation (Shi et al. 2022), recent work suggests that bone marrow located within the skull calvaria, skull base, and vertebrae plays roles in meningeal and CNS immunity (Mazzitelli et al. 2023; Cugurra et al. 2021).

In mice, the cribriform plate region of the skull is a dynamic site which facilitates the drainage of CSF, waste, antigen, and immune cells from the intracranial space into the meningeal-nasal lymphatic system (Hsu et al. 2019, 2022; Choi et al. 2025; C. J. Laaker et al. 2025; Hong et al. 2026; C. J. Laaker et al. 2026; C. Laaker et al. 2026; Yoon et al. 2024; H. Jin et al. 2025; Ma et al. 2019; Decker et al. 2022). In mice, the cribriform plate bone marrow (cpBM), is positioned dorsal-rostral to the olfactory bulb and flanks the portion of the environment between skull and nasal regions (C. J. Laaker et al. 2025). Recently, our lab has identified that skull channels link some portions of cribriform plate bone marrow to the cerebrospinal fluid (CSF) interfacing environment near the olfactory bulb, cribriform lymphatics, and olfactory nerve bundles (C. J. Laaker et al. 2025; Hsu et al. 2019; C. J. Laaker et al. 2026). Furthermore, in these skull channels, we observed CSF1R-GFP+, CD45+, and CD4+ immune cells, suggesting active immune trafficking between environments (Laaker et al., 2025). During EAE, the cpBM displays markers of elevated myelopoiesis, similar to femur bone marrow (Laaker et al., 2025), but it remains unclear whether immune cells derived from the cpBM region and broader nasal environment have unique characteristics that allow them to differentially regulate immunity such as immunosuppression (Hsu et al. 2022; Shi et al. 2022; C. J. Laaker et al. 2025). Thus, uncovering the role cpBM region could reveal new therapeutic targets and strategies for inducing immune tolerance in neuroinflammatory disease. .

In this study, we sought to understand three main questions. (1) How does the cpBM region change during EAE? (2) Is the cpBM region distinct compared to systemic BM sites in the femur during EAE? (3) Does this region have distinct immunoregulatory abilities on pathogenic T cells? To answer these questions we utilized flow, IHC and single-cell RNA sequencing (scRNAseq) and identified the immune composition of cpBM region isolates. During EAE, we found that there is an expansion of myeloid subsets collected from location, particularly semi-mature and mature Ly6G+ granulocytes with an upregulation of inflammatory and migratory genes (Monaghan et al. 2019; Wang et al. 2024). During EAE, the cpBM region displayed evidence of neutrophils with transcription factors related to late stage tissue-infiltration and immunoregulation. Additionally, to assess the functional impact of these shifts, we performed flow cytometry, and co-culture experiments comparing cpBM region to cell derived from femur bone marrow. During EAE the cpBM region samples had an expanded pool of hematopoietic stem cells (HSCs) and Tregs. Interestingly, cpBM region cultures had a suppressive effect on myelin specific CD4⁺ T cells which differentiate into IL-10+ Tregs, in contrast to femur marrow, which promoted pro-inflammatory IFN-γ+ effector T cells. Finally, we show that the cpBM region contains an elevated proportion of developing and immature B cells in EAE. Together, these findings support that cribriform plate bone marrow region undergoes immune remodeling during peak EAE, ultimately with the broader environment having a distinct ability to influence T cells. Our results highlight cpBM region as a unique immune site at the CNS border, with implications for understanding immune surveillance, drainage, and immunoregulation during neuroinflammation.

## Results

### Cribriform plate bone marrow is integrated via vascular skull channels with the perineural environment at the cribriform plate

We previously identified the cpBM skull region to be an interconnected bone marrow environment near CSF interfacing olfactory nerve bundles in mice (C. J. Laaker et al. 2025). Indeed, in vivo intravascular labeling of osseous channels with Osteosense EX680 and CD31 antibody confirms the cpBM has vascular connections in IHC of decalcified sections of the perineural space of healthy mouse olfactory nerve bundles (**Fig 1A-C; Supplemental Movie 1**). Intra-cisterna magna injection of Evan’s Blue dye directly into the CSF of EAE mice confirmed the cribriform bone marrow regions near nerve bundles contained measurable levels of injected dye (**Fig 1D-G**).

**Figure 1.**
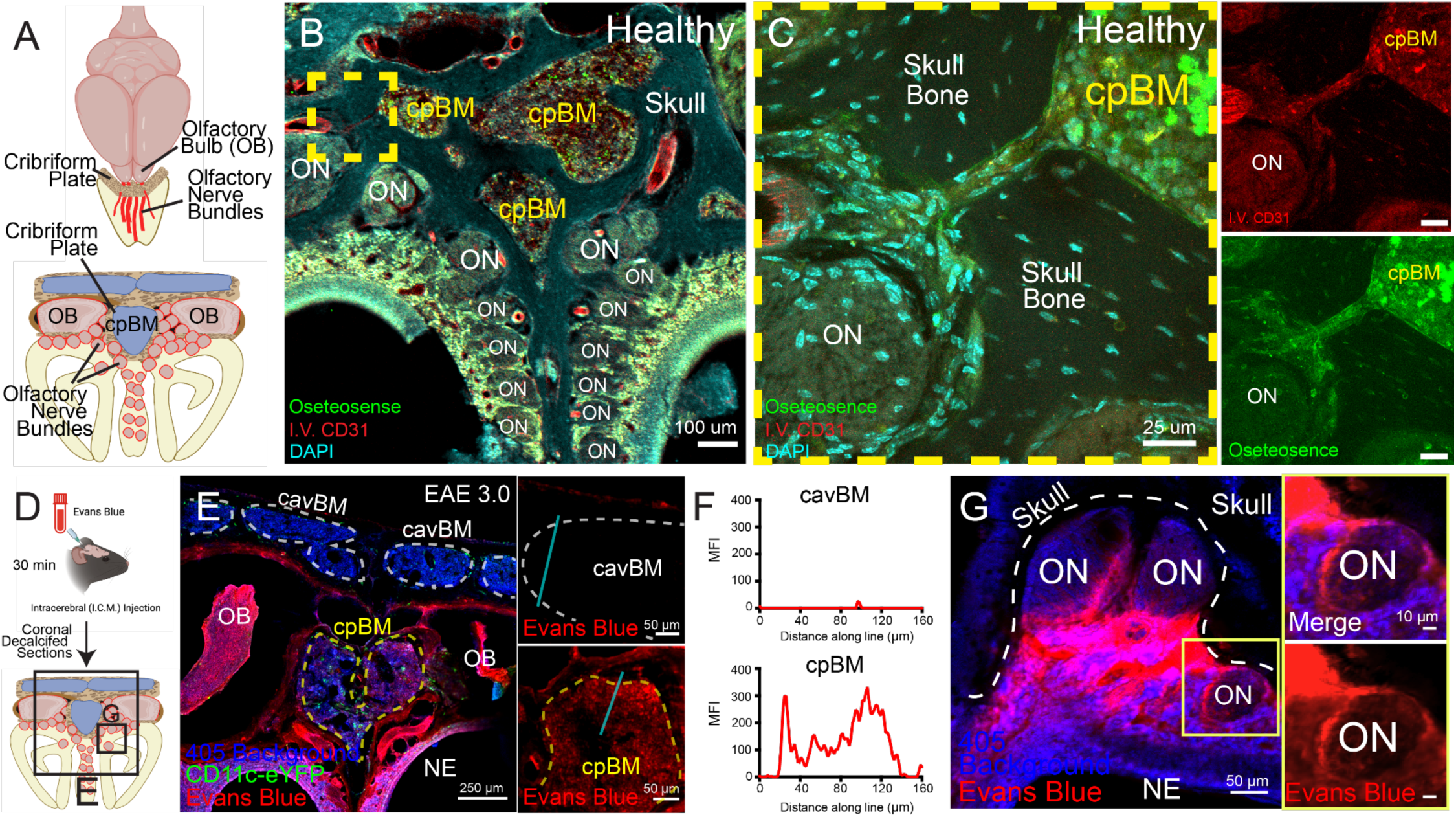
Cribriform Plate Bone Marrow has perineural vascular channels and displays access to ICM injected dye. **(A):** Live healthy mice were retro-orbitally injected with OsteoSense 680EX and CD31 PE antibody 24 hours before tissue collection. Whole head cribriform plate sections were decalcified and imaged with a confocal microscope. Representative image shows a large pool of cribriform plate bone marrow (cpBM) in close association with several olfactory nerve bundles (ON) passing through cribriform plate foramina. n = 3 **(B-C):** Zoom of perineural skull channel (yellow dotted square) is positive for CD31 (red) blood vessels and boundaries of skull channel are positive for osteosense (green), confirming these are osseous conduits with vascular connections. **(D-G):** EAE mice were injected with 5ul of Evans Blue Dye (Red) and euthanized 30min later (**D**). Whole heads were decalcified and analyzed for the presence of dye (**E-F**). Evans blue accumulates perineurally around olfactory nerve bundles (ON) (**G**).

### Cribriform plate bone marrow region immune cells have a unique expressional profile during EAE

To understand how CNS autoimmunity reshapes the cpBM region in mice, we performed single-cell RNA sequencing on cpBM region sample preparations from healthy and EAE mice, and inflamed systemic bone marrow from EAE femurs (**Fig. 2A-C)**. Importantly, our cpBM skull isolates are highly enriched with the local bone marrow, but due to the interwoven anatomy of the sampled region (**Figure 1B**) contain portions of the surrounding foramina environment, olfactory nerve bundles, and underlying mucosal environment (**Supplemental Figure 1**). Indeed, IHC confirmed pockets of EAE 3.0 cpBM had high levels of Ly6G+ cells in CSF1R-GFP mice, highlighting dense myeloid cell populations near the olfactory bulb and olfactory nerve bundles (**Fig. 2A-B**). For sequencing we isolated cpbM regions from healthy and EAE 3.0 mice, and additionally Femurs from EAE 3.0 mice. UMAP clustering of scRNAseq data revealed distinct shifts in cellular composition between healthy and EAE conditions in the cpBM region, with notable expansion of semi-mature and mature Ly6G+ populations in EAE cpBM (**Fig. 2D-E**). Quantitative comparison showed increased representation of myeloid cell subsets including cycling and maturing Ly6G+ cells, accompanied by a relative reduction in B cells, progenitor populations, and T cells (**Fig. 2E-F**). These cell proportions closely resembled slight elevations of lymphoid populations in cpBM vs Femur observed in flow cytometry (**Fig. 2C**), but still supports an overall shift toward myeloid lineage during EAE as reported previously (C. J. Laaker et al. 2025).

**Figure 2.**
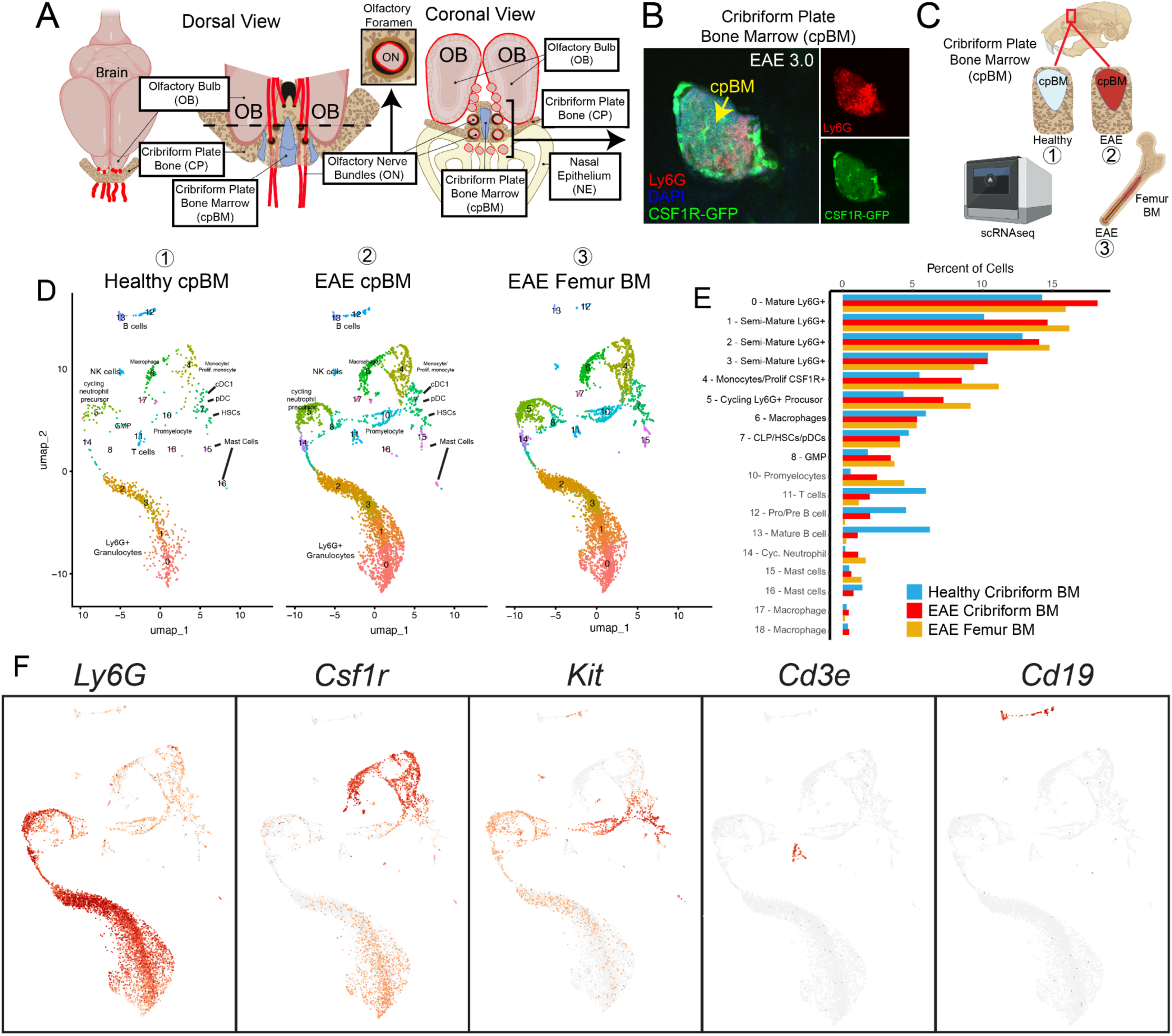
Experimental setup and scRNAseq cell cluster identities from cpBM and Femur BM. **(A)** Cartoon scheme of the dorsal and coronal view of the olfactory bulb (OB) region highlighting the midline cribriform plate (CP) and olfactory nerve bundles (ON) traversing the bone toward the nasal epithelium (NE). **(B)** Representative image of cpBM from a EAE 3.0 CSF1R-GFP mouse stained with Ly6G. **(C)** Healthy and EAE 3.0 cribriform plate (CNS-adjacent bone) and EAE 3.0 femur (systemic bone) were excised from C57/BL6 mice. Cribriform plate bone marrow cells were extracted by gently crushing the cribriform plate bone, while femur bone marrow cells were isolated by injecting RPMI 1640 medium through the femoral medullary cavity. **(D)** UMAP plot showing successful clustering of cpBM cells from scRNA-seq with dominant myeloid populations. Bar graph displaying the relative proportions of the 18 identified cpBM clusters from scRNA-seq analysis. Cluster 9 was neuronal debris and dead cells and was excluded from analysis. **(E)** Cell proportions of each cluster in each experimental groups, note the increase in myeloid populations during EAE **(F)** Common cell markers used for cluster identification and Heatmap output from Loupe Browser 8.0

Overall EAE induced a broad inflammatory signature in the cpBM on Ly6G, monocyte, and macrophage cluster subsets when compared to Healthy (**Figure 3A**). Transcriptomic profiling of the largest cluster (Ly6G+) revealed significant upregulation of inflammatory and migratory genes in cpBM compared to femur during EAE, including *Ifitm1*, *Lrg1*, and *Cxcr2* (**Fig. 3B**). In order to determine how the cpBM is distinct from peripheral bone marrow sites, we compared EAE cpBM vs EAE Femur BM (**Figure 3C**). During EAE the cpBM had higher levels of several genes suggestive of mature tissue infiltrating and/or immunosuppressive neutrophils including *Cebpb*, *Pim1*, *Nfkbia*, *Dusp1* (**Fig. 3D**) (Volberding et al. 2021; Alshetaiwi et al. 2020; S. Jin et al. 2024). Of note, *Cebpb* (C/EBPβ) is understood to be a “master regulator” of generation and is a key driver of MDSC immunosuppressive activity (Fultang et al. 2020; He et al. 2025). For example, in tumor-infiltrating myeloid cells C/EBPβ promotes a negative influence on T Cell activation (Marigo et al. 2010).

**Figure 3.**
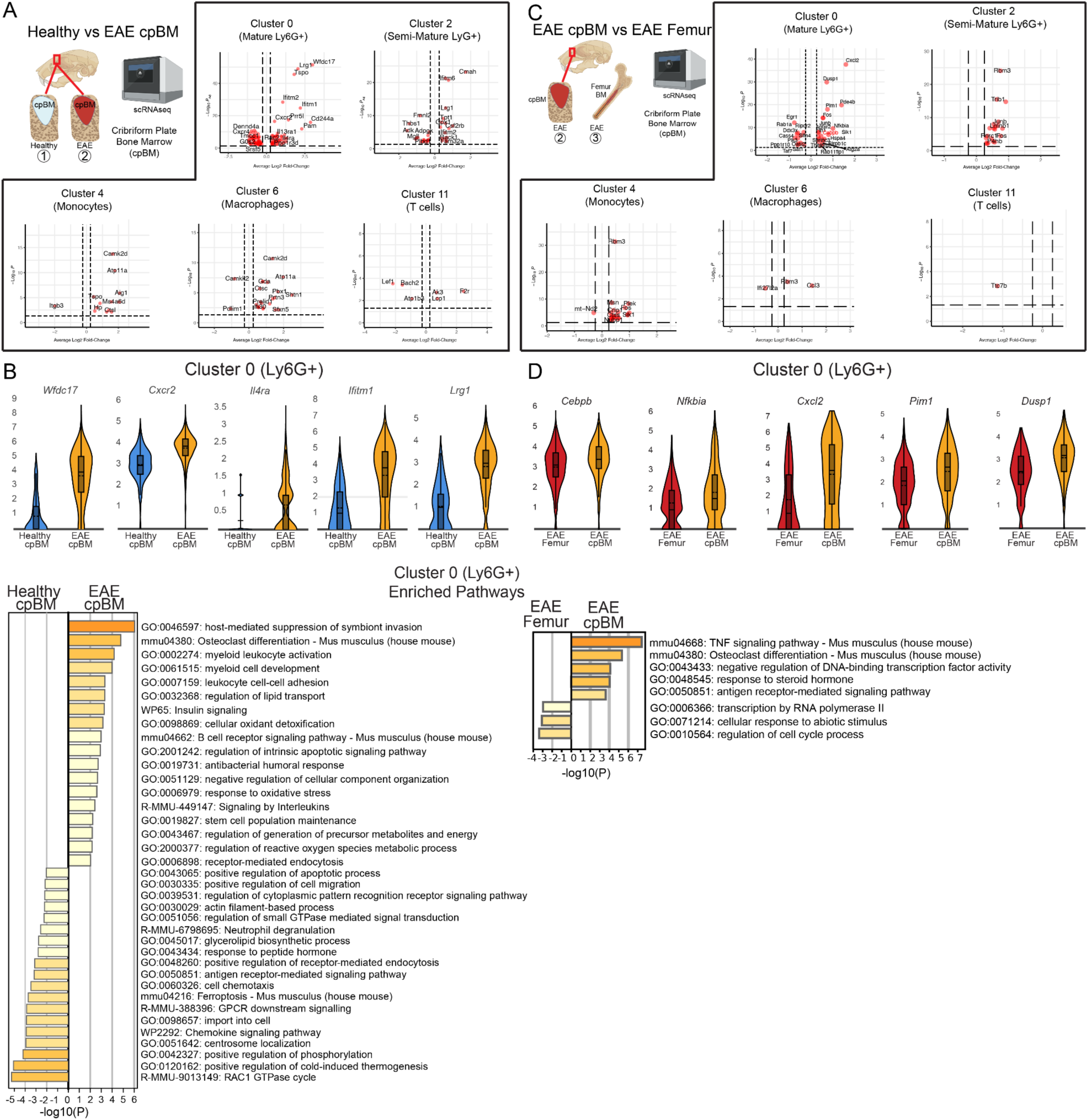
EAE shifts cpBM toward unique myelopoietic state. **(A-B)** Analysis of healthy vs EAE cpBM. Volcano plots comparing cpBM across inflammatory conditions in key clusters. Cutoff of an adjusted p-value < 0.01 to determine DEGs **(A)**. Violin plots showing upregulated genes in Cluster 0 (Mature Ly6G+) immune cells from EAE cpBM compared to Healthy cpBM **(B)**. **(C-D)** Analysis of EAE cpBM vs Femur bone marrow. Volcano plots comparing inflamed EAE cpBM vs Femur bone marrow. Cutoff of an adjusted p-value < 0.01 to determine DEGs **(C)**. Violin plots showing upregulated genes in Cluster 0 (Mature Ly6G+) immune cells from EAE cpBM compared to EAE Femur BM **(D)**.

To understand if T cell populations (*Cd3e*+ cells) differ across bone marrow niches, we analyzed differentially expressed genes from each sample. Healthy cpBM T cells exhibited a naïve/stem-like state, while EAE cpBM T cells displayed a distinct profile characterized by elevated proliferation and immune regulation, differing even from EAE femur bone marrow T cells (**Figure 4A-C**). This shift in EAE cpBM was marked by increased expression of cell cycle genes (Ki67, Cenpa, Cks2) and regulatory molecules often associated with Treg phenotype (Il2ra, Nrp1, Ctla4, Lgals3) (Setoguchi et al. 2005; Walker 2013; Yadav et al. 2012; Ocklenburg et al. 2006; Jaroušek et al. 2022). Pathway analysis further highlighted enrichment for cell division, Th17 differentiation, and chemotaxis in EAE cpBM T cells, contrasting the more homeostatic and survival pathways in healthy cpBM (**Figure 4B-C**).

**Figure 4.**
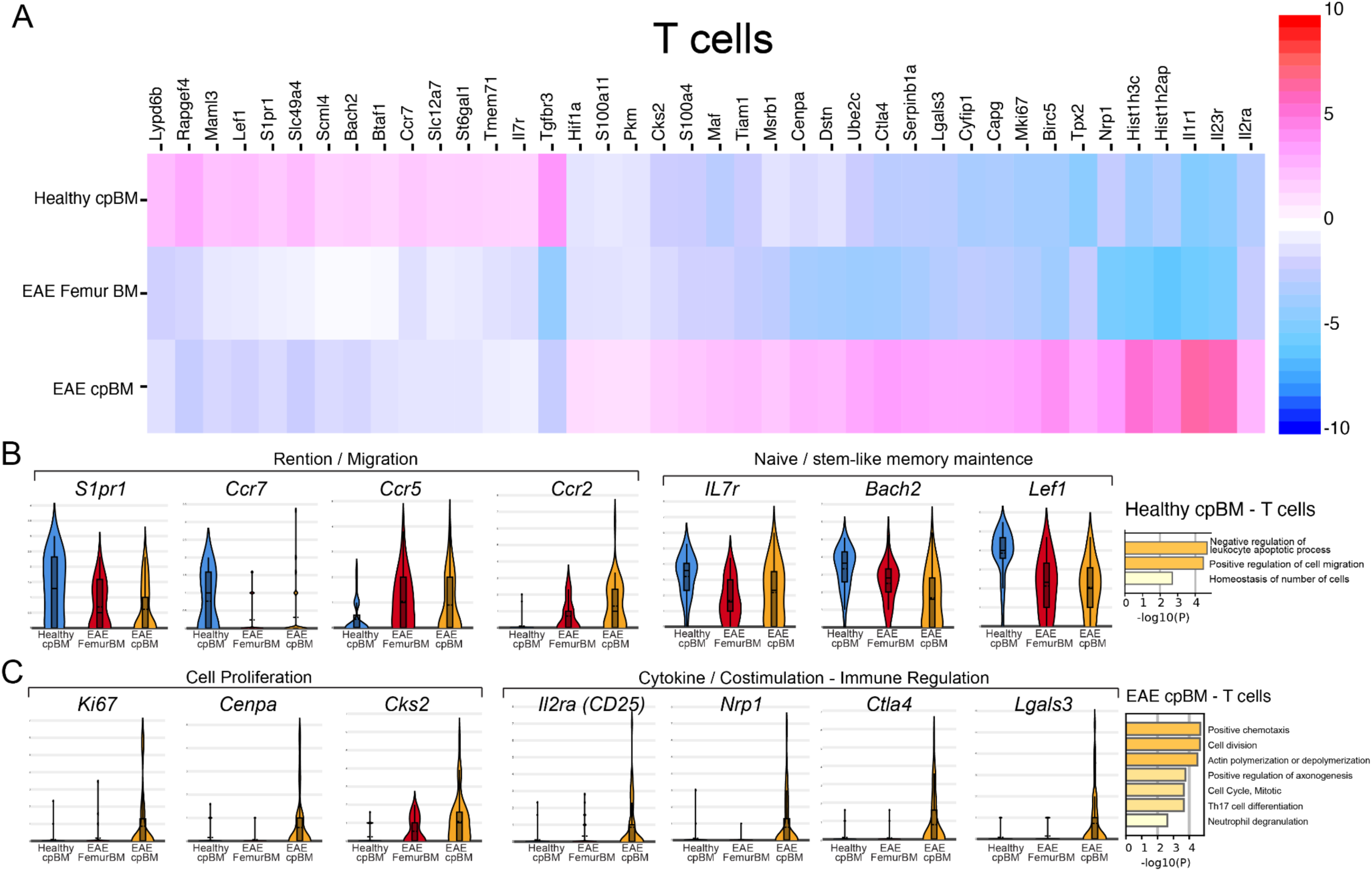
Distinct transcriptional programs of T cells in cpBM during EAE. **(A) :** Heatmap (top) shows relative gene expression (row-normalized) of T cells (*Cd3e*+ cells) from healthy cpBM, EAE femur BM, and EAE cpBM. **(B-C):** Violin plots illustrate relevant expression of genes grouped by function: migration/retention (*S1pr1, Ccr7*), naïve/stem-like maintenance (*IL7r, Bach2, Lef1*), proliferation (*Mki67, Cenpa, Cks2*), and immune regulation (*Il2ra/CD25, Nrp1, Ctla4, Lgals3*).

These findings confirm that while cpBM undergoes a myeloid shift during EAE similar to femur, its transcriptional profile suggests heightened MDSC potential (Alshetaiwi et al. 2020). Furthermore, our evidence supports that T cells isolated from cpBM have a unique transcriptional profile suggestive of either Treg / Th17 differentiation when compared to both healthy cpBM and EAE Femur BM.

### Cribriform plate bone marrow region has elevated HSC populations

Given the cpBM has a unique position between mucosal layers, nerve bundles, and the CNS, we next investigated how EAE influences the hematopoietic activity in the cribriform plate bone marrow region versus femur (systemic) bone of EAE 3.0 mice (**Fig. 2A**). To test this we collected samples from the cribriform plate bone marrow region (**Fig. 2A, Supplemental Figure 1**), and analyzed via flow cytometry to identify HSCs and mature immune cell subsets (**Fig. 2B**).

To validate our sequencing data we measured the frequencies of myeloid populations (neutrophils, dendritic cells, and monocytes) and lymphoid populations (CD4, CD8, B cells) in cribriform and femur BM isolates during EAE (**Fig 5A**). Similar to our sequencing data, EAE femur and CP isolates have similar levels of myeloid cells (**Fig. 5B**). Interestingly, cpBM isolates had elevated levels of CD4 T cells and B cells compared to femur (**Fig. 5C**). Additionally, HSCs (Lin⁻ c-Kit⁺ Sca1⁺ CD48⁻ CD150⁺) were significantly elevated in the cpBM isolates (**Fig. 5D**). Together this data supports similar myeloid cell composition across femur and cpBM, but cpBM isolates contain elevated levels of lymphoid and HSC population.

**Figure 5.**
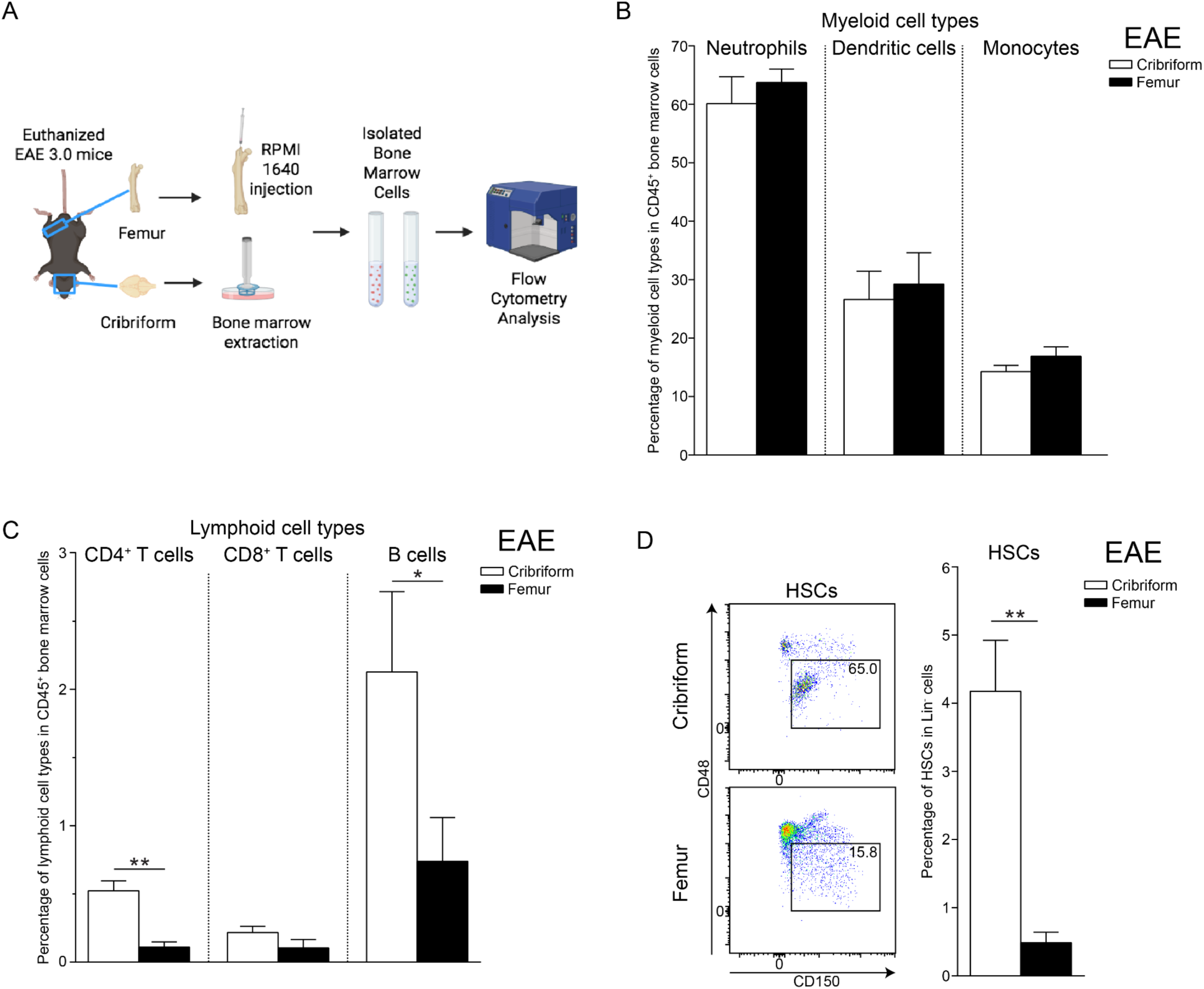
Cribriform Plate Bone Marrow Region has elevated HSC and lymphoid populations compared to femur during EAE. **(A)** Experimental scheme. Cribriform plate (CNS-adjacent bone) and femur (systemic bone) were excised from EAE 3.0 mice. Cribriform plate bone marrow cells were extracted by gently crushing the cribriform plate bone, while femur bone marrow cells were isolated by injecting RPMI 1640 medium through the femoral medullary cavity. **(B)** Percentage of indicated myeloid cell types in CD45^+^ cribriform plate and femur bone marrow of EAE 3.0 mice. Mean ± SEM. n = 5 mice per group. **(C)** Percentage of indicated lymphoid cell types in CD45^+^ cribriform plate and femur bone marrow of EAE 3.0 mice. Mean ± SEM. n = 5 mice per group. Unpaired two-tailed Student’s *t*-test. \**p* < 0.05; \*\**p* < 0.01. **(D)** Representative gating strategy for flow cytometry analysis of cellular composition of cribriform plate and femur bone marrow of EAE 3.0 mice. Percentage of HSCs in Lin^-^ cribriform plate and femur bone marrow of EAE 3.0 mice. Mean ± SEM. n = 3 mice per group. Unpaired two-tailed Student’s *t*-test. \*\**p* < 0.01.

### Weakened Myelin-antigen specific T cell activation in cribriform plate bone marrow region at peak EAE

With the observation that the cpBM has suppressive potential and T cells might differ across, we sought to determine whether cpBM differentially modulates the activation of CNS-antigen specific T cells during neuroinflammation. To do this we established ex vivo co-cultures of MOG-specific 2D2 CD4⁺ T cells with bone marrow cultures isolated from the cribriform plate and femur of EAE 3.0 mice (**Fig. 6A**). CD4⁺ T cells from 2D2 TCR-MOG transgenic mice were co-cultured with marrow cells each niche, and T cell activation was assessed by expression of CD69 at 24 hours, CD40L at 48 hours, and LFA-1 at 72 hours (**Fig. 6A**). Flow cytometry gating identified 2D2 CD4⁺ T cells (CD4⁺, TCR Vβ11⁺) within co-cultures (**Fig. 6B**). Representative plots showed reduced expression of activation markers in 2D2 T cells co-cultured with cribriform plate marrow compared to those co-cultured with femur marrow (**Fig. 6C**). Quantification confirmed significantly lower frequencies of CD69⁺, CD40L⁺, and LFA-1⁺ 2D2 T cells in the cpBM across all time points (**Fig. 6D**), indicating a locally immunosuppressive niche that dampens antigen specific T cell activation in CNS-adjacent bone marrow.

**Figure 6.**
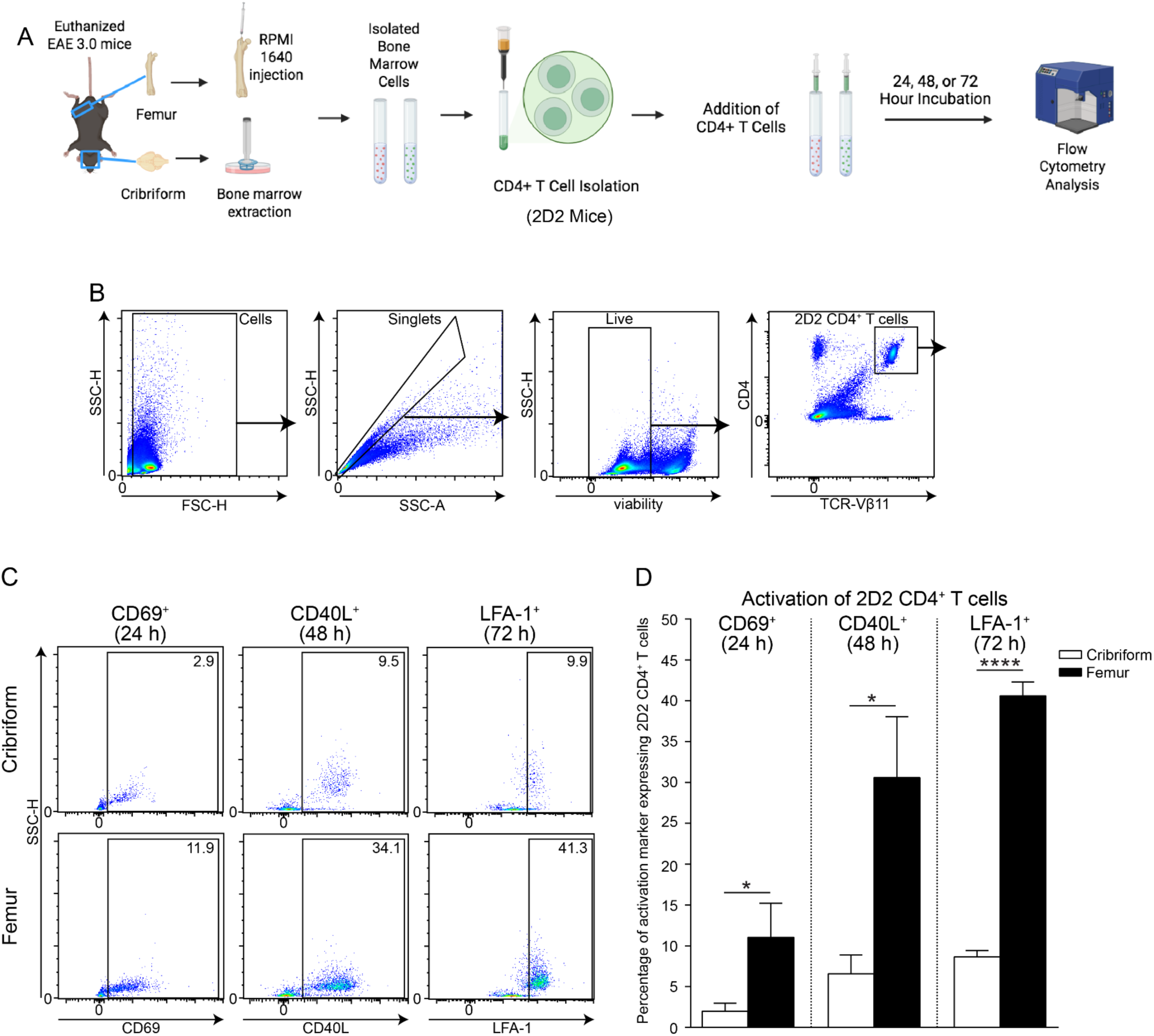
CNS Autoimmunity Dampens T Cell Activation in Cribriform Plate Bone Marrow Region. **(A)** Experimental scheme. Cribriform plate (CNS-adjacent bone) and femur (systemic bone) were excised from EAE 3.0 mice. Cribriform plate bone marrow cells were extracted by gently crushing the cribriform plate bone, while femur bone marrow cells were isolated by injecting RPMI 1640 medium through the femoral medullary cavity. Bone marrow cell suspensions were transferred to separate wells of a sterile 96-well plate. CNS-antigen (MOG) specific 2D2 CD4^+^ T cells were isolated from the spleen of a healthy 2D2 TCR^MOG^ transgenic mouse using a CD4^+^ T cell isolation kit via magnetic-activated cell sorting (MACS). 2 × 10^4 2D2 CD4^+^ T cells were added to each well containing the various bone marrow cells to initiate 2D2 CD4^+^ T cell – bone marrow cell co-cultures. Following incubation for 24, 48, or 72 hours, respectively, the co-cultures underwent staining for flow cytometry to analyze the effects of different bone marrows in the induction of activation marker expression on 2D2 CD4^+^ T cells: CD69 after 24 hours, CD40L after 48 hours, and LFA-1 after 72 hours. **(B)** Representative gating strategy for flow cytometry analysis of 2D2 CD4^+^ T cells (CD4^+^, TCR Vβ11^+^) in 2D2 CD4^+^ T cell – bone marrow cell co-cultures of cribriform plate and femur bone marrow of EAE 3.0 mice. **(C)** Representative flow cytometry plots show the percentage of expression of activation markers CD69 after 24 hours, CD40L after 48 hours, and LFA-1 after 72 hours on 2D2 CD4^+^ T cells that had been co-cultured with cribriform plate or femur bone marrow cells of EAE 3.0 mice. **(D)** Percentage of activation marker CD69, CD40L, or LFA-1 expressing 2D2 CD4^+^ T cells after having been co-cultured with cribriform plate or femur bone marrow cells of EAE 3.0 mice for 24, 48, or 72 hours, respectively. Mean ± SEM. n = 3 mice per group. Unpaired two-tailed Student’s *t*-test. \**p* < 0.05; \*\*\*\**p* < 0.0001.

### Cribriform Plate Bone Marrow region favors regulatory T Cell differentiation during neuroinflammation

To examine how CNS-adjacent and systemic bone marrow environments influence the differentiation and polarization of CNS-antigen specific T cells during neuroinflammation, 2D2 CD4⁺ T cells were transferred *in vivo* into EAE 3.0 mice and analyzed 72 hours later from Cribriform plate and femur bone marrow (**Fig. 7A**). Flow cytometry gating identified donor 2D2 T cells (CD4⁺, TCR Vβ11⁺) within each marrow compartment (**Fig. 7B**). Representative plots showed that the cribriform plate environment supported a higher frequency of regulatory T cells (Tregs; FoxP3⁺) and tissue-memory T cells (CD44⁺CD62L⁺), whereas the femur promoted differentiation toward effector T cells (CD44⁺CD62L⁻) (**Fig. 7C**). Quantification confirmed a significant increase of Tregs in cpBM region cultures, whereas effector T cells were induced by femur bone marrow (**Fig. 7D**). Cytokine analysis further revealed that 2D2 tissue memory T cells in the cribriform plate marrow were more likely to produce the anti-inflammatory cytokine IL-10, while those in the femur produced more IFN-γ (**Fig. 7E-F**). These findings were shared across in effector subsets also, with elevated IL-10⁺ cells in the cribriform plate and IFN-γ⁺ cells in the femur (**Fig. 7G-H**). Together, these results suggest that the cpBM region promotes regulatory T cells, while systemic marrow favors a more pro-inflammatory phenotype.

**Figure 7.**
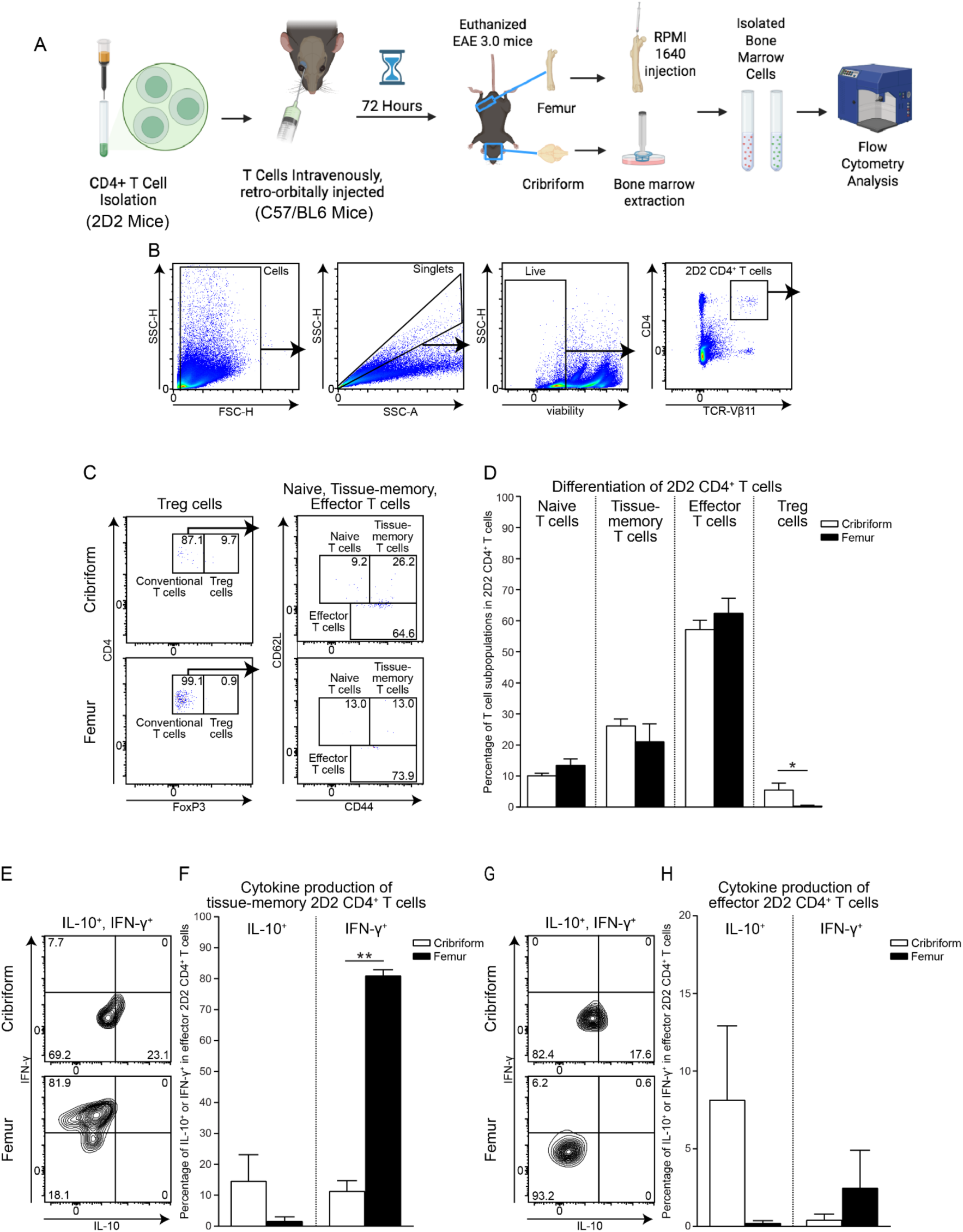
Cribriform Plate Bone Marrow Region Supports Anti-Inflammatory T Cell Fate in CNS Autoimmunity. **(A) :** Experimental scheme. CNS-antigen (MOG) specific 2D2 CD4^+^ T cells were isolated from the spleen of a healthy 2D2 TCR^MOG^ transgenic mouse using a CD4^+^ T cell isolation kit via magnetic-activated cell sorting (MACS). EAE 3.0 mice were intravenously, retro-orbitally injected with 1.5 x 10^7^ 2D2 CD4^+^ T cells/mouse. Following 72 hours, the cribriform plate (CNS-adjacent bone) and femur (systemic bone) were excised from the EAE 3.0 mice. Cribriform plate bone marrow cells were extracted by gently crushing the cribriform plate bone, while femur bone marrow cells were isolated by injecting RPMI 1640 medium through the femoral medullary cavity. The bone marrow cell suspensions underwent staining for flow cytometry to analyze the effects of different bone marrow niches in the induction of T cell differentiation from naïve T cells (CD44^-^CD62L^+^) toward tissue-memory T cells (CD44^+^CD62L^+^), effector T cells (CD44^+^CD62L^-^), and Treg (FoxP3^+^) cells; as well as in the induction of T cell polarization toward pro-inflammatory (IFN-γ) or anti-inflammatory (IL-10) cytokine producer tissue-memory T cells (CD44^+^CD62L^+^) and effector T cells (CD44^+^CD62L^-^). **(B)** Representative gating strategy for flow cytometry analysis of 2D2 CD4^+^ T cells (CD4^+^, TCR Vβ11^+^) in bone marrow cell suspensions of cribriform plate and femur bone marrow of EAE 3.0 mice that had been injected with 2D2 CD4^+^ T cells 72 hours prior to the experiment. **(C)** Representative flow cytometry plots show the percentage of naïve T cells (CD44^-^CD62L^+^), tissue-memory T cells (CD44^+^CD62L^+^), effector T cells (CD44^+^CD62L^-^), and Treg (FoxP3^+^) cells within 2D2 CD4^+^ T cells in cribriform plate and femur bone marrow of EAE 3.0 mice that had been injected with 2D2 CD4^+^ T cells 72 hours prior to the experiment. **(D)** Percentage of naïve T cells (CD44^-^CD62L^+^), tissue-memory T cells (CD44^+^CD62L^+^), effector T cells (CD44^+^CD62L^-^), and Treg (FoxP3^+^) cells within 2D2 CD4^+^ T cells in cribriform plate and femur bone marrow of EAE 3.0 mice that had been injected with 2D2 CD4^+^ T cells 72 hours prior to the experiment. Mean ± SEM. n = 3 mice per group. Unpaired two-tailed Student’s *t*-test. \**p* < 0.05. **(E-G)** Representative flow cytometry plots show the percentage of IFN-γ (pro-inflammatory) and IL-10 (anti-inflammatory) cytokine producer tissue-memory T cells (CD44^+^CD62L^+^) and effector T cells (CD44^+^CD62L^-^) within 2D2 CD4^+^ T cells in cribriform plate and femur bone marrow of EAE 3.0 mice that had been injected with 2D2 CD4^+^ T cells 72 hours prior to the experiment. **(F-H)** Percentage of IFN-γ (pro-inflammatory) and IL-10 (anti-inflammatory) cytokine producer tissue-memory T cells (CD44^+^CD62L^+^) and effector T cells (CD44^+^CD62L^-^) within 2D2 CD4^+^ T cells in cribriform plate and femur bone marrow of EAE 3.0 mice that had been injected with 2D2 CD4^+^ T cells 72 hours prior to the experiment. Mean ± SEM. n = 2-3 mice per group. Unpaired two-tailed Student’s *t*-test. \*\**p* < 0.01.

### Cribriform Plate Bone Marrow Region is enriched for immature B Cells in EAE

Finally, since MDSCs and suppressive bone marrow niches can inhibit B cell proliferation, function, and maturity (Jaufmann et al. 2020; Crook et al. 2015) we assessed B cell dynamics in cpBM versus femur bone marrow during EAE neuroinflammation (**Fig. 8A**). Flow cytometry gating was used to identify CD19⁺B220⁺ B cells within the CD45⁺ bone marrow (**Fig. 8B-C**). Quantification confirmed a significant increase in the frequency of B cells in cpBM region (**Fig. 8D**). Further characterization of B cell developmental stages showed that the cribriform plate contained a higher proportion of pro B cells (IgM⁻IgD⁻) and immature B cells (IgM⁺IgD⁻), while the femur marrow was more enriched in mature B cells (IgD⁺) (**Fig. 8E-F**). Together these data indicate that CNS-adjacent bone marrow in the cpBM supports larger, but more immature B cells during EAE when compared to Femur.

**Figure 8.**
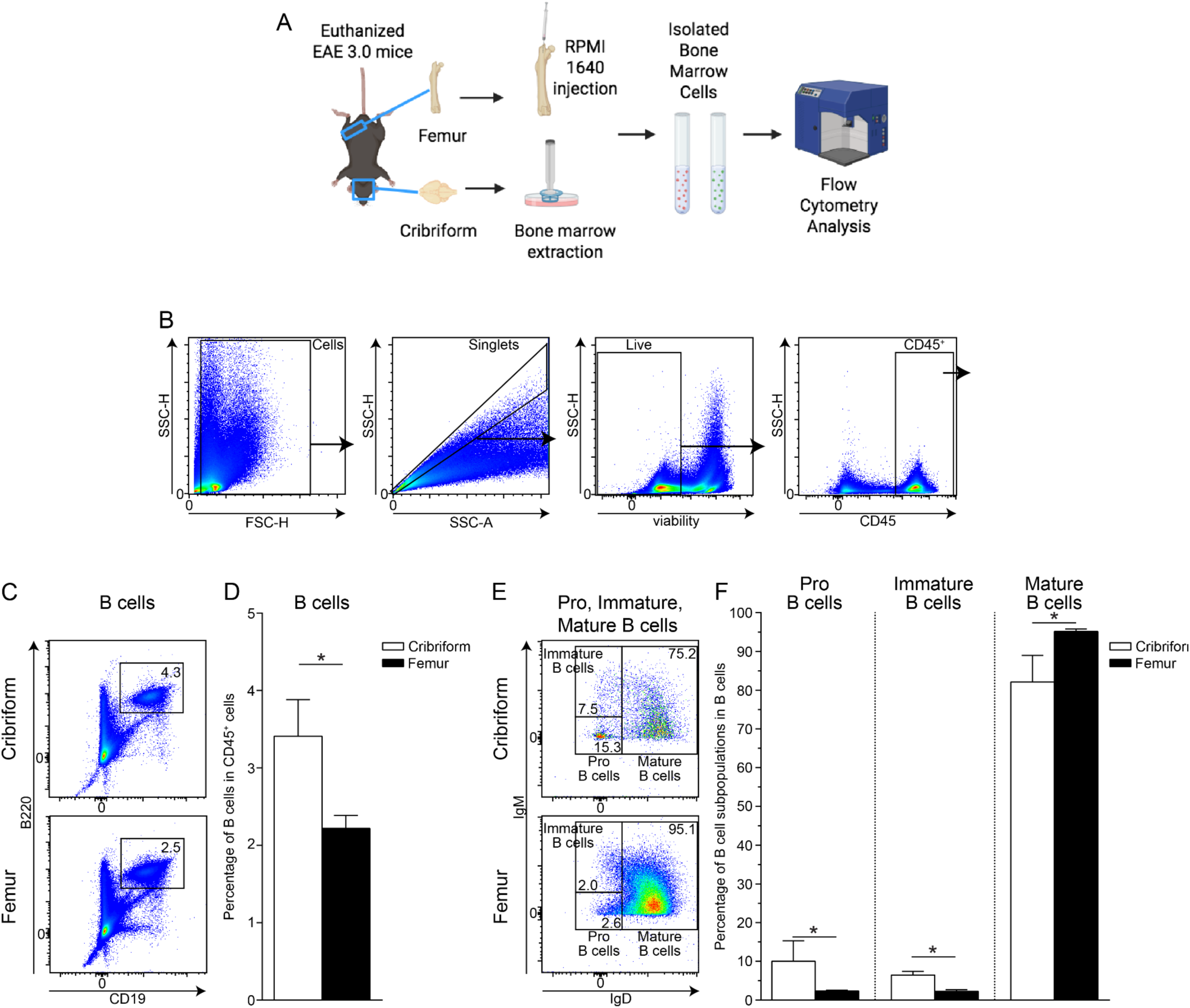
Cribriform Plate Bone Marrow Region Enriches for Immature B Cells in Autoimmunity. **(A)** Experimental scheme. Cribriform plate (CNS-adjacent bone) and femur (systemic bone) were excised from EAE 3.0 mice. Cribriform plate bone marrow cells were extracted by gently crushing the cribriform plate bone, while femur bone marrow cells were isolated by injecting RPMI 1640 medium through the femoral medullary cavity. **(B)** Representative gating strategy for flow cytometry analysis of B cells in bone marrow cell suspensions of cribriform plate and femur bone marrow of EAE 3.0 mice. **(C)** Representative flow cytometry plots show the percentage of B cells (CD19^+^, B220^+^) in CD45^+^ cribriform plate and femur bone marrow of EAE 3.0 mice. **(D)** Percentage of B cells in CD45^+^ cribriform plate and femur bone marrow of EAE 3.0 mice. Mean ± SEM. n = 3 mice per group. Unpaired two-tailed Student’s *t*-test. \**p* < 0.05. **(E)** Representative flow cytometry plots show the percentage of B cell developmental stages: pro B cells (IgM^-^, IgD^-^), immature B cells (IgM^+^, IgD^-^), and mature B cells (IgD^+^) within B cells in cribriform plate and femur bone marrow of EAE 3.0 mice. **(F)** Percentage of B cell developmental stages: pro B cells (IgM^-^, IgD^-^), immature B cells (IgM^+^, IgD^-^), and mature B cells (IgD^+^) within B cells in cribriform plate and femur bone marrow of EAE 3.0 mice. Mean ± SEM. n = 2-3 mice per group. Unpaired two-tailed Student’s *t*-test. \**p* < 0.05.

## Discussion

Our findings show that the cribriform plate bone marrow (cpBM) region, a CNS-adjacent hematopoietic niche, undergoes distinct immunological remodeling during CNS autoimmunity. Unlike femur bone marrow, the cpBM displays several unique features in cellular composition, with expansion of hematopoietic stem cells, myeloid cell subsets, and accumulation of developmentally immature B cells. Functionally, our data suggests cpBM region can contribute as a locally immunosuppressive hub to the activation of CNS-antigen specific CD4⁺ T cells; skewing differentiation toward regulatory phenotypes at peak EAE. In femur marrow however, cultures promoted effector T cell responses. Single-cell transcriptomics further revealed enhanced myeloid skewing in EAE cpBM, particularly among Ly6G+ subsets, there was an upregulation of inflammatory, migratory, and MDSC related gene expression like *Cebpb* (Alshetaiwi et al. 2020; Fultang et al. 2020) and *Pim1 (*Volberding et al. 2021; S. Jin et al. 2024). Additionally, T cells isolated from the EAE cpBM region had gene signatures related to Treg and/or Th17 phenotype. Together, these data suggest cpBM is a distinct immunological compartment potentially serving as a local reservoir to regulate immune activity near the cribriform - olfactory bulb interface.

In the context of the CNS, the skull and vertebra bone marrow are emerging as important sources of local immunity. In mice, the cpBM region is indicated to be a unique site of immune activity (Choi et al. 2025; Hsu et al. 2022, 2019; C. J. Laaker et al. 2025; C. Laaker et al. 2026; C. J. Laaker et al. 2026). Adjacent to several olfactory nerve bundles and lymphatic vessels, cpBM is implicated to be exposed to CSF-derived cues and antigens (C. J. Laaker et al. 2025; Hsu et al. 2019). This study demonstrates that cpBM region exerts a unique immunological profile during EAE, highlighting it as not only a source of immune cells but a modifier of T cell responses, shifting toward tolerance rather than inflammation. There are several limitations to this study, for one the exact cell type and mechanism in the bone marrow environment contributing to cpBM immunosuppression is still not fully elucidated. Influences of T cell inhibition pathways can be complex with contributions due to shifts in local immunometabolism (Arginase-1, etc) or checkpoint molecules (PD-L1, VISTA). Furthermore, while our bone marrow preparations were consistent, highly concentrated with myeloid cells and our scRNAseq supported highly overlapping signature between femur and cpBM region preps, it is ultimately highly challenging to isolate foramina interfacing skull bone marrow regions like the cpBM, and some non-bone marrow cells including nerves and mucosal tissue likely influence our cpBM enriched samples. Thus these findings should be viewed as an investigation of the broader cpBM niche. Future studies will investigate alternative techniques such as spatial transcriptomics to specifically validate these findings.

In conclusion, our study shows that the cpBM region undergoes a myeloid remodeling during EAE which has distinct attributes when compared to systemic sites. Flow cytometry and scRNA sequencing of the cpBM skull region points to an expansion of HSCs, an immune profile skewing toward MDSCs, and increased B cells and CD4⁺ T cell proportions. In contrast, femur marrow in EAE favors a more classical myelopoiesis and contributes toward effector T cell polarization. These findings suggest that cpBM may serve a more inhibitory function in neuroinflammation. Functionally, this study shows that cpBM region derived cells suppress the activation of myelin-antigen specific 2D2 CD4⁺ T cells. When 2D2 CD4⁺ T cells were co-cultured with cells isolated from the cpBM region, they exhibited reduced expression of activation markers (CD69, CD40L, LFA-1) and differentiated into IL-10 producing regulatory T cells. By contrast, T cells cultured with femur marrow had higher activation and IFN-γ production, characteristic of a pro-inflammatory effector type. These observations indicate the cpBM region is a locally immunosuppressive niche, possibly contributing to CNS immune privilege through promotion of antigen-specific tolerance. Additionally, these results support that bone marrow niches across the body can exert distinct and competing impacts on lymphoid cells during autoimmunity and raise the possibility of targeting skull bone marrow regions for therapeutic purposes.

## Methods

### Animals

Female wild-type (WT) C57BL/6J mice (Strain: 000664), 2D2 TCR^MOG^ transgenic mice (expressing T cell receptors (TCRs) specific for myelin oligodendrocyte glycoprotein (MOG), Strain: 006912), and CSF1R-EGFP “MacGreen” transgenic mice (expressing enhanced green fluorescent protein (EGFP) in colony stimulating factor 1 receptor (CSF1R) myeloid cells, Strain: 018549) were purchased from The Jackson Laboratory. Female mice aged 8–12 weeks were used for all EAE experiments, along with corresponding age- and sex-matched controls. All experiments were performed in compliance with the guidelines from the National Institutes of Health and the University of Wisconsin–Madison Institutional Animal Care and Use Committee.

### EAE Induction

Experimental autoimmune encephalomyelitis (EAE) was induced in female mice aged 8 - 12 weeks through subcutaneous immunization with 100 μg of MOG_35–55_ peptide (Cat: MOG3555-P-5, Genemed Synthesis Inc.) emulsified in Complete Freund’s Adjuvant (CFA, Cat: F5881, Sigma-Aldrich) at the dorsal region between the shoulder blades. Following immunization, 200 ng of pertussis toxin (PT, Cat: NC9675592, List Biological Laboratories Inc) was administered intraperitoneally at 0- and 2-days post-immunization. The onset of clinical symptoms was observed between 8- and 12-days post-immunization. Clinical scores were monitored daily, applying the following scoring system: 0 = no clinical symptoms, 1 = limp or flaccid tail, 2 = partial hind limb paralysis, 3 = complete hind limb paralysis, 4 = quadriplegia, and 5 = moribund. Intermediate scores were assigned when appropriate. Mice that reached an EAE clinical score of 3.0 by day 15-18 post-immunization were selected for all subsequent experiments conducted on EAE mice.

### Bone Marrow Cell Isolation

Healthy and EAE 3.0 mice were euthanized under terminal anesthesia with isoflurane and perfused transcardially with PBS (Cat: 21-040-CM, Corning). Following decapitation, incisions were made to remove the skullcap, leading to the exposure of the brain. The olfactory bulbs were dissected out and placed on a 40 µm cell strainer set in a Petri dish containing Roswell Park Memorial Institute (RPMI) 1640 medium (Cat: 10-040-CV, Thermo Fisher Scientific). Meningeal cells were mechanically dissociated with the plunger of a sterile syringe. The cribriform plate bone was carefully excised, rinsed with PBS, and placed on a 40 µm cell strainer set in a RPMI 1640 medium containing Petri dish. Cribriform plate bone marrow cells were extracted by gently crushing the cribriform plate bone with the plunger of a sterile syringe. The femur was also removed, and the femur bone marrow cells were flushed out by injecting RPMI 1640 medium through the femoral medullary cavity using a sterile syringe. The cells were washed 3x in 900 µl RPMI 1640 medium (1200 rpm, 10 min, 4 °C).

### T Cell Activation Assay

Equal portions of cribriform plate and femur bone marrow cell suspensions extracted from EAE 3.0 mice were transferred to separate wells of a sterile round bottom tissue culture-treated 96-well plate (Cat: 353077, Corning) in 200 µl/well cell culture medium consisting of RPMI-1640 medium supplemented with 10% heat-inactivated fetal bovine serum (FBS, Cat: A5256701, Thermo Fisher Scientific), 2-mercaptoethanol (Cat: M3148, Sigma-Aldrich), and penicillin/streptomycin. For positive control, equal portions of femur bone marrow cell suspensions extracted from EAE 3.0 mice were seeded under identical conditions. 2D2 CD4^+^ T cells were isolated from the spleen of a healthy 2D2 TCR^MOG^ transgenic mouse using a CD4^+^ T cell isolation kit (Cat: 130-104-454, Miltenyi Biotec) via magnetic-activated cell sorting (MACS) according to the manufacturer’s instructions. Then, 2 × 10^4 2D2 CD4^+^ T cells were added to each well containing the various bone marrow cells to initiate 2D2 CD4^+^ T cell – bone marrow cell co-cultures. For negative control, 2 × 10^4 2D2 CD4^+^ T cells were seeded in 200 µl cell culture medium to separate wells in the absence of bone marrow cells. The positive control co-cultures were supplemented with 100 µg/ml MOG_35–55_ peptide (Cat: MOG3555-P-5, Genemed Synthesis Inc.) as well. The co-cultures were incubated for 24, 48, or 72 hours, respectively, at 37 °C and 5% CO_2_; then underwent staining for flow cytometry to analyze the effects of different bone marrow niches in the induction of activation marker expression on 2D2 CD4^+^ T cells: CD69 after 24 hours (Cat: 553236, BD Biosciences), CD40L (CD154) after 48 hours (Cat: 157011, BioLegend), and LFA-1 (CD11a/CD18) after 72 hours (Cat: 25-0111-82, Thermo Fisher Scientific). Detailed information about all antibodies applied in this study is provided in Table 1.

**Table 1.**
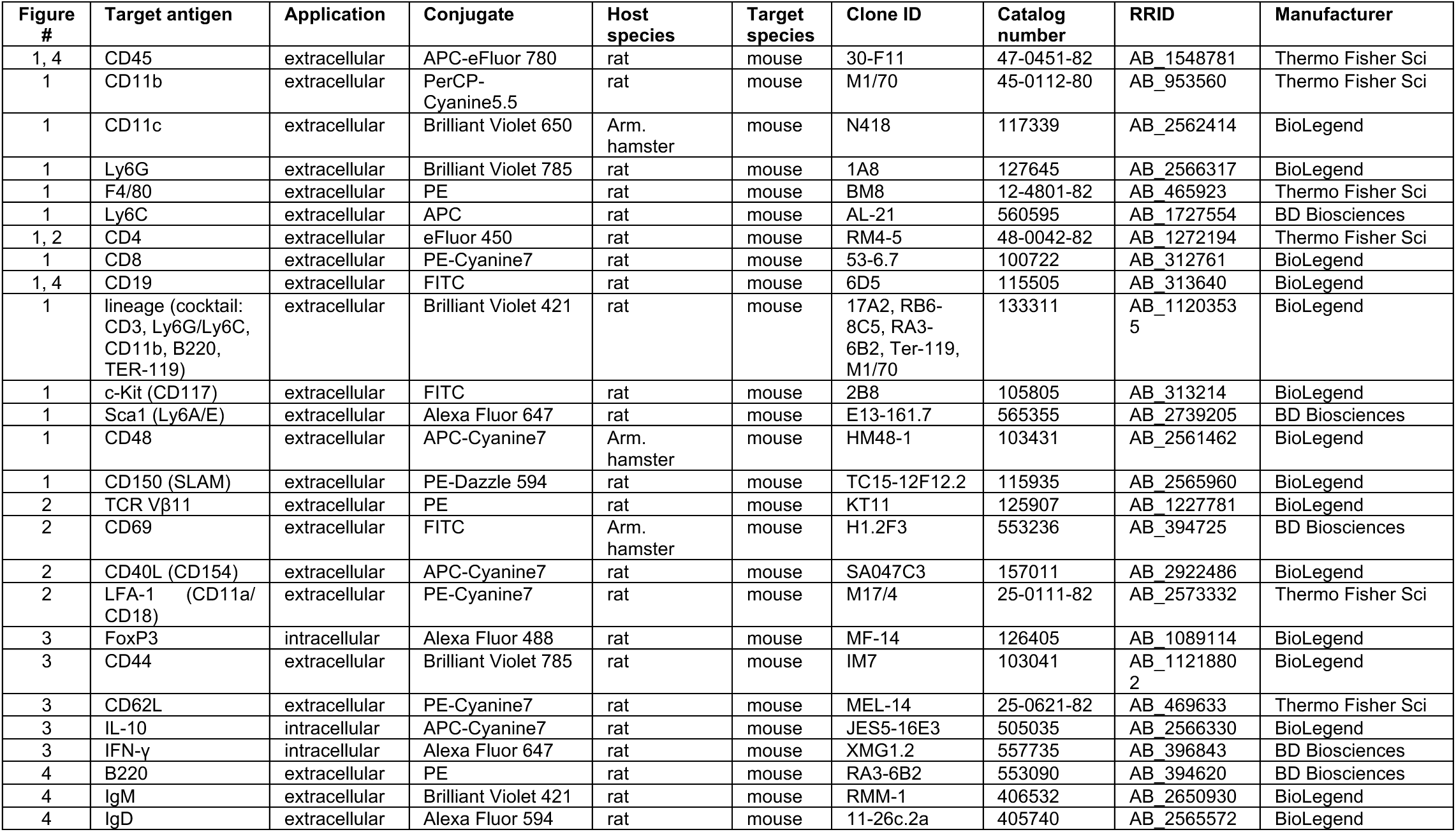
Antibodies.

**Table 2.**
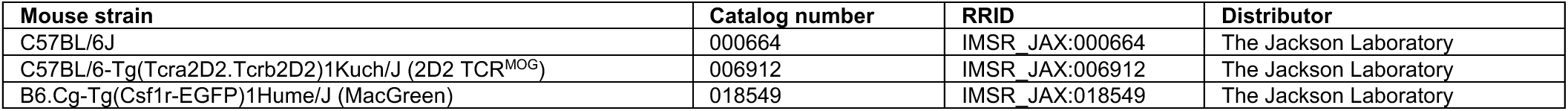
Mouse strains.

| Mouse strain | Catalog number | RRID | Distributor |
| --- | --- | --- | --- |
| C57BL/6J | 000664 | IMSR_JAX:000664 | The Jackson Laboratory |
| C57BL/6-Tg(Tcrα2D2.Tcrβ2D2)1Kuch/J (2D2 TCR<sup>MOG</sup>) | 006912 | IMSR_JAX:006912 | The Jackson Laboratory |
| B6.Cg-Tg(Csf1r-EGFP)1Hume/J (MacGreen) | 018549 | IMSR_JAX:018549 | The Jackson Laboratory |

**Table 3.** Reagents, peptides.

| Reagents, peptides | Catalog number | Manufacturer |
| --- | --- | --- |
| MOG<sub>35-55</sub> peptide | MOG3555-P-5 | Genemed Synthesis Inc. |
| Complete Freund's Adjuvant | F5881 | Sigma-Aldrich |
| pertussis toxin | NC9675592 | Thermo Fisher Scientific |
| isoflurane | NDC 59399-106-01 | Akorn Inc. |
| PBS | 21-040-CM | Corning |
| paraformaldehyde | 15714-S | Electron Microscopy Sciences |
| EDTA | 02-152-52191 | Thermo Fisher Scientific |
| sucrose | S24065 | Research Products International |
| Tissue-Tek O.C.T. Compound | 4583 | Sakura Finetek USA |
| RPMI 1640 medium | 10-040-CV | Thermo Fisher Scientific |
| FBS | A5256701 | Thermo Fisher Scientific |
| 2-mercaptoethanol | M3148 | Sigma-Aldrich |
| GolgiStop Protein Transport Inhibitor | 51-2092KZ | BD Biosciences |
| Ghost Violet 540 Viability Dye | 13-0879-T100 | Tonbo Biosciences |
| BSA | A3294 | Sigma-Aldrich |

**Table 4.** Commercial kits.

| Commercial assay | Catalog number | Manufacturer |
| --- | --- | --- |
| CD4<sup>+</sup> T cell isolation kit | 130-104-454 | Miltenyi Biotec |
| Foxp3/Transcription Factor Staining Buffer Set | 00-5523-00 | Thermo Fisher Scientific |
| ArC Amine Reactive Compensation Bead Kit | A10346 | Thermo Fisher Scientific |
| UltraComp eBeads Compensation Beads | 01-2222-42 | Thermo Fisher Scientific |

**Table 5.** Software.

| Software | Manufacturer |
| --- | --- |
| FlowJo v10 | BD Biosciences |
| GraphPad Prism 6 | GraphPad Software |

### T Cell Differentiation and Cytokine Production Assay

The wells of a sterile round bottom tissue culture-treated 96-well plate (Cat: 353077, Corning) were coated with CD3 specific antibody (Cat: 16-0031-86, Thermo Fisher Scientific) at a concentration of 4 µg/ml in 50 µl sterile PBS (Cat: 21-040-CM, Corning), and incubated overnight at 4 °C. Prior to cell addition, the wells were further supplemented with CD28 specific antibody (Cat: 553295, BD Biosciences) at 4 µg/ml concentration in additional 50 µl sterile PBS. Then, equal portions of cribriform plate and femur bone marrow cell suspensions extracted from EAE 3.0 mice were transferred to separate wells of the plate in 100 µl/well cell culture medium (ingredients described above). In this way, the cells were stimulated with 1 µg/ml final concentration of both plate-bound CD3 and CD28 specific antibodies. For positive control, equal portions of femur bone marrow cell suspensions extracted from EAE 3.0 mice were seeded under identical conditions. 2D2 CD4^+^ T cells were isolated from the spleen of a healthy 2D2 TCR^MOG^ transgenic mouse using a CD4^+^ T cell isolation kit (Cat: 130-104-454, Miltenyi Biotec) via magnetic-activated cell sorting (MACS) according to the manufacturer’s instructions. Then, 2 × 10^4 2D2 CD4^+^ T cells were added to each well containing the various bone marrow cells to initiate 2D2 CD4^+^ T cell – bone marrow cell co-cultures. For negative control, 2 × 10^4 2D2 CD4^+^ T cells were seeded in 200 µl cell culture medium to separate wells in the absence of bone marrow cells. The positive control co-cultures were supplemented with 100 µg/ml MOG_35–55_ peptide (Cat: MOG3555-P-5, Genemed Synthesis Inc.) as well. The co-cultures were maintained for 6 days at 37 °C and 5% CO_2_, with careful replacement of the cell culture media every 24 hours. On day 6, the cell culture medium in each well was replaced with GolgiStop Protein Transport Inhibitor (Cat: 51-2092KZ, BD Biosciences) diluted 1:1500 in 200 µl/well cell culture medium, according to the manufacturer’s instructions. The co-cultures were incubated for additional 5 hours at 37 °C and 5% CO_2_, then underwent staining for flow cytometry to analyze the effects of different bone marrow niches in the induction of 2D2 CD4^+^ T cell differentiation from naïve (CD44^-^CD62L^+^) toward T CM (central memory, CD44^+^CD62L^+^), T EM (effector memory, CD44^+^CD62L^-^) (103041, BioLegend; Cat: 25-0621-82, Thermo Fisher Scientific) or Treg (regulatory, FoxP3^+^) cells (Cat: 126405, BioLegend); as well as in the induction of IL-10 (Cat: 505035, BioLegend), IL-17 (Cat: 45-7177-82, Thermo Fisher Scientific), and IFN-γ (Cat: 557735, BD Biosciences) cytokine production in 2D2 CD4^+^ T cells. Detailed information about all antibodies applied in this study is provided in Table 1.

### Flow Cytometry

For the flow cytometric analysis of freshly isolated bone marrow and meningeal cells, as well as of 2D2 CD4^+^ T cell – bone marrow cell co-cultures, the cells were first washed 3x in PBS (Cat: 21-040-CM, Corning) (1200 rpm, 10 min, 4 °C), and then stained with Ghost Violet 540 Viability Dye (Cat: 13-0879-T100, Tonbo Biosciences) in PBS for 30 minutes at 4 °C, following the manufacturer’s protocol to distinguish viable from non-viable cells. After washing 3x in 900 µl PBS containing 1% BSA (bovine serum albumin, Cat: A3294, Sigma-Aldrich) (1200 rpm, 10 min, 4 °C), the cells were stained extracellularly with fluorescent dye conjugated antibodies in 200 µl PBS containing 1% BSA for 1 hour at 4 °C, and then were washed 3x in 900 µl PBS containing 1% BSA (1200 rpm, 10 min, 4 °C). Additionally, the cells underwent fixation, permeabilization, and intracellular staining with fluorescent dye conjugated antibodies for 30 min at 4 °C, by the application of the Foxp3/Transcription Factor Staining Buffer Set (Cat: 00-5523-00, Thermo Fisher Scientific) and following the manufacturer’s instructions. Finally, the cells were washed 3x in 900 µl PBS containing 1% BSA (1200 rpm, 10 min, 4 °C) and resuspended in 200 µl PBS containing 1% BSA. Flow cytometry analysis was performed on a Northern Lights flow cytometer (Cytek Biosciences). Compensation was performed using the ArC Amine Reactive Compensation Bead Kit (Cat: A10346, Thermo Fisher Scientific) and UltraComp eBeads Compensation Beads (Cat: 01-2222-42, Thermo Fisher Scientific). Flow cytometric data were analyzed using FlowJo v10 (BD Biosciences). Detailed information about all antibodies applied in this study is provided in Table 1.

### Statistics

An unpaired Student’s *t*-test was used to compare results between two different groups. All statistical analyses were performed using GraphPad Prism 6 (GraphPad Software). Data are presented as mean ± SEM. Significance is displayed as NS (not significant), \**P* < 0.05, \*\**P* < 0.01, \*\*\**P* < 0.001, or \*\*\*\**P* < 0.0001.

### scRNAseq Tissue Harvest and Cell Isolation

Following a 3-minute incubation, mice were euthanized without perfusion. The cribriform plate bone, containing bone marrow, was dissected, rinsed with phosphate-buffered saline (PBS), and placed on a 40 μm cell strainer within a Petri dish containing RPMI 1640 medium (Thermo Fisher Scientific). The bone was mechanically disrupted using the plunger from a sterile syringe to release bone marrow cells into the surrounding medium. Separately, the femur was harvested, and the bone marrow was flushed out by injecting RPMI medium through the bone shaft using a syringe. The cell suspensions from both tissues were transferred to 15 ml Falcon tubes and centrifuged at 1,250 rpm for 10 minutes at 4 °C.

Cribriform plate bone marrow cells from three EAE 3.0 mice were pooled into a single “EAE cribriform” sample. Femur bone marrow from the same three mice was pooled into an “EAE femur” sample. The healthy cribriform sample was obtained from a single age-matched healthy control mouse. All samples were resuspended in 5 ml of 1× RBC Lysis Buffer (eBioscience, Cat# 00-4333-57, Lot# 2362015), incubated with constant shaking for 5 minutes, and centrifuged at 1,250 rpm for 10 minutes at 4 °C. Pellets were washed and resuspended in PBS and filtered through 40 μm cell strainers into sterile Eppendorf tubes. Each sample was incubated with 2 μl Ghost UV 450 viability dye (Tonbo Biosciences, Cat# 13-0868-T100, Lot# D0868083018133) for 10 minutes at 4 °C. Following viability staining, the following reagents were added per sample: 10 μl fetal bovine serum (FBS), 5 μl anti-mouse CD16/CD32 FcR-blocking reagent (BD Pharmingen, Clone 2.4G2, Cat# 553142, Lot# 7248907), 5 μl TotalSeq-B oligo-conjugated sample-specific hashtag antibody: EAE cribriform: Hashtag 11 (anti-CD45 and anti-MHC-I, BioLegend Cat# 113901, Lot# B385027), EAE femur: Hashtag 12 (anti-CD45 and anti-MHC-I, BioLegend Cat# 113903, Lot# B385029), Healthy cribriform: Hashtag 13 (anti-CD45 and anti-MHC-I, BioLegend Cat# 113905, Lot# B385030). Samples were incubated for 30 minutes at 4 °C, then washed with 1 ml PBS + 5% FBS, centrifuged at 1,250 rpm for 10 minutes at 4 °C, and resuspended in 300 μl PBS + 5% FBS. FACS sorting: single cells were sorted based on viability dye exclusion and FSC/SSC parameters. Sorted cell numbers were as follows: 60,000 from the healthy cribriform sample, 100,000 from the EAE cribriform sample, and 100,000 from the EAE femur sample. All sorted populations were pooled into a single tube for downstream single-cell RNA sequencing (scRNA-seq), with individual sample identities preserved via oligo-conjugated hashtag antibodies.

### scRNA Sequencing

Sorted bone marrow was then provided to the Biotechnology Core facility at the University of Wisconsin Madison for single cell sequencing using the 10x Genomics Chromium Single Cell Gene Expression Assay. Cells were sorted and provided to the Biotechnology Core for scRNA-seq. Samples were then washed and resuspended with a FACS buffer (1% BSA + 1x PBS). Cells were loaded onto a Chromium Controller to generate a single cell + barcoded gel-bead emulsion. Libraries were prepped with the 10X Genomics 3’ reagents kit (v3 chemistry). The target recovery rate was 3,000 cells with a targeted read depth of 76,000 per cell. Cells were sequenced on the NovaSeq S1 100-cycle flow cell in collaboration with the University of Wisconsin Biotechnology Center (UWBC) DNA Sequencing Facility. Data were analyzed by the UW Bioinformatics Resource Center. Experimental data were demultiplexed using the Cell Ranger Single Cell Software Suite, mkfastq command wrapped around Illumina’s bcl2fastq. The Miseq balancing run was quality controlled using calculations based on UMI tools (Smith et al. 2017). Sample libraries were balanced for the number of estimated reads per cell and run on an Illumina NovaSeq system. Cell Ranger software was then used to perform demultiplexing, alignment, filtering, barcode counting, UMI counting, and gene expression estimation for each sample according to the 10x Genomics documentation. Barcodes containing unusually high numbers of detected transcripts indicative of a doublet signature were excluded, and cells that co-express marker genes of non relevant cell types were also excluded. The resulting data were then analyzed and explored using the Loupe Cell Browser software and the R packages clusterProlifer and/or monacle3 after excluding cells that made it through the FACS-enrichment. Cells have > 200 genes expressed, Cells have < 3,500 genes expressed, Cells have < 5% mitochondrial genes expressed.

### Intravascular and Osteosense labeling

2 nmol of IVISense Osteo 680 (NEV10020EX) was I.P. injected into 8 week old healthy mice 24 hours before tissue collection. In the same mouse, to label blood vasculature, 2.5 μg of CD31 antibody conjugated to PE (BioLegend, catalog no. 102507) was injected retro orbitally in 200 μl of PBS (Anderson et al. 2014). Mice were perfused and whole heads were decalcified according to conditions below.

### Histology

Mice underwent terminal anesthesia with isoflurane, followed by transcardial perfusion with 0.1 M PBS and subsequent perfusion with 4% paraformaldehyde (PFA) in 0.1 M PBS. Post-perfusion, mice were decapitated, and the skin encompassing the head was carefully removed using forceps and scissors to separate it from the muscle and ear canal. The entire heads were fixed in 4% PFA in 0.1 M PBS overnight. Subsequently, the heads underwent decalcification in 14% EDTA in 0.1 M PBS for 7 days, followed by cryoprotection in 30% sucrose in 0.1 M PBS for 3 days. Fresh 14% EDTA was replaced daily during decalcification. The decalcified mouse heads were then embedded in Tissue-Tek OCT compound, frozen on dry ice, and stored at −80°C. Cryostat-cut sections measuring 60 μm in thickness were obtained using a Leica CM1800 cryostat, mounted on Superfrost Plus microscope slides, and stored at −80°C. For serial sections, each section was labeled and stored in accordance with its slice position.

---

**Supplementary Figure 1.**
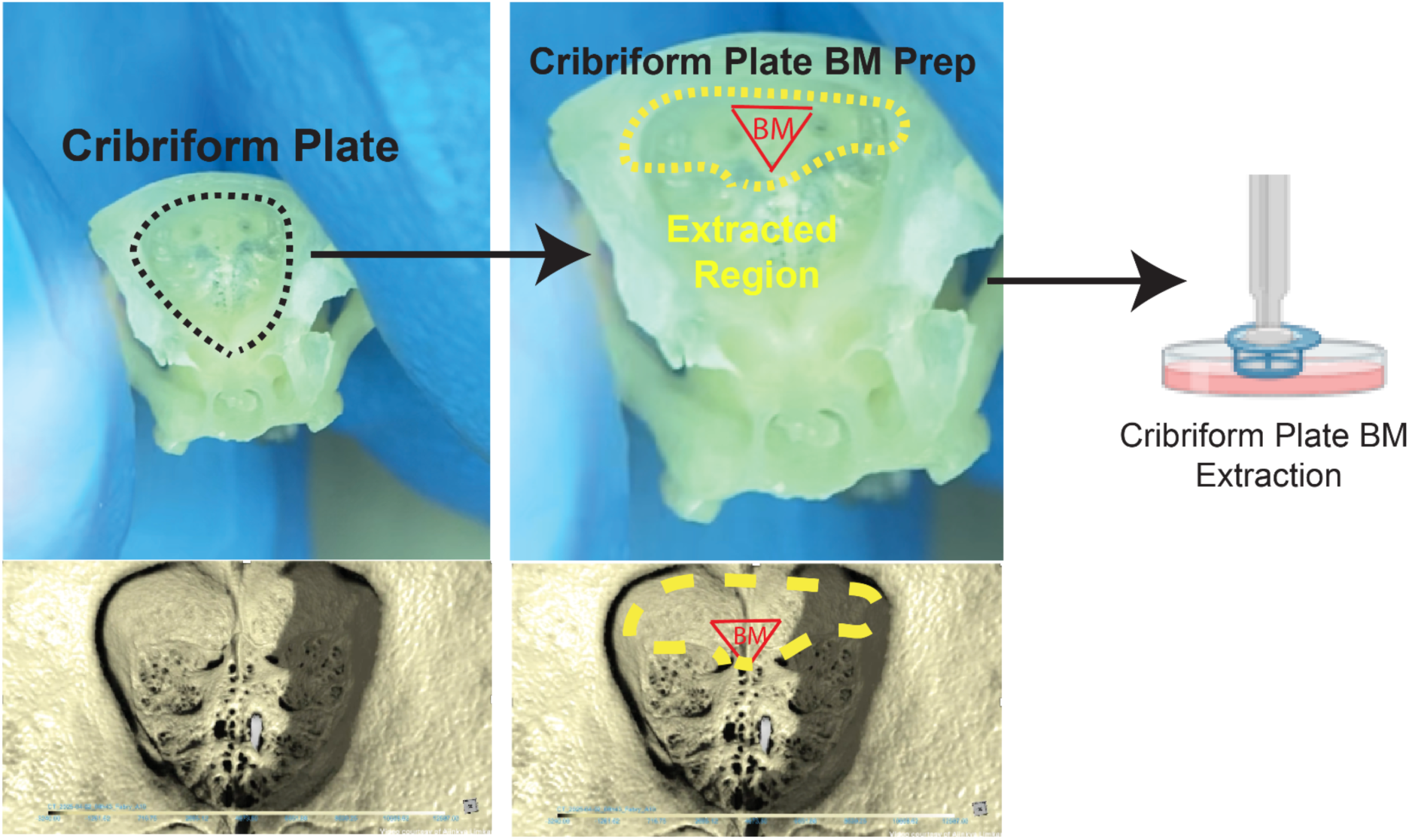
Representative images showing isolated cpBM regions. Samples were washed with PBS prior to dissociation. Samples were enriched for cpBM but contained a broader cpBM region environment as well. CT image provided by Ajinkya R Limar and Sophia Vrba.

**Supplementary Figure 2.**
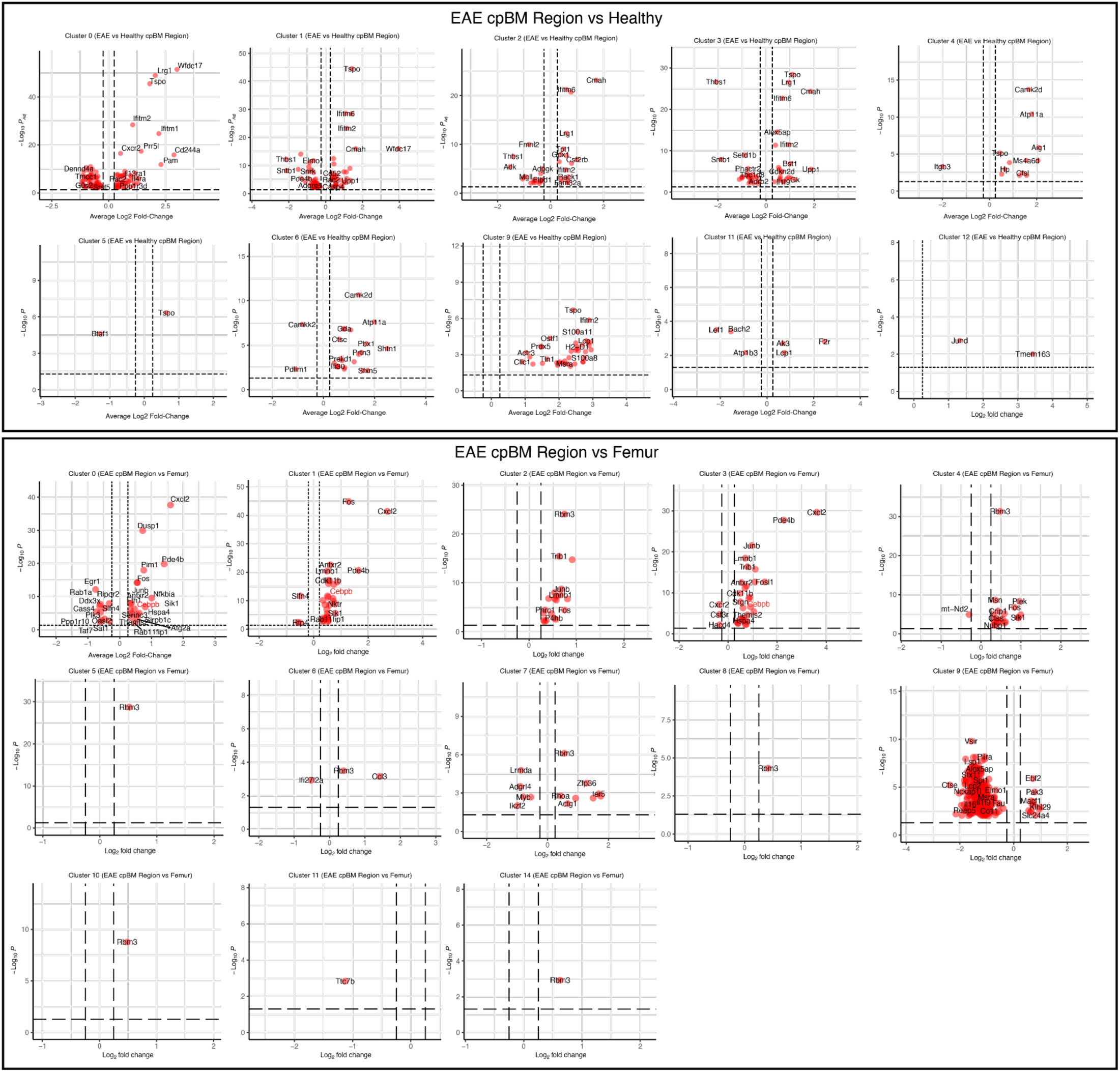
Volcano plots for clusters with significant differences between EAE cpBM vs Healthy cpBM or EAE feumur BM. Top plots show EAE vs Healthy cpBM comparisons, while bottom shows EAE cpBM region vs EAE femur.

